# Representing Multiple Observed Actions in the Motor System

**DOI:** 10.1101/387704

**Authors:** Emiel Cracco, Christian Keysers, Amanda Clauwaert, Marcel Brass

**Author notes:** Correspondence concerning this article should be addressed to Emiel Cracco, Department of Experimental Psychology, Ghent University, Henri Dunantlaan 2, B-9000.

## Abstract

There is now converging evidence that others’ actions are represented in the motor system. However, social cognition requires us to represent not only the actions but also the interactions of others. To do so, it is imperative that the motor system can represent multiple observed actions. The current fMRI study investigated whether this is possible by measuring brain activity from 29 participants while they observed two right hands performing sign language gestures. Three key results were obtained. First, brain activity in the premotor and parietal motor cortex was stronger when two hands performed two different gestures than when one hand performed a single gesture. Second, both individual observed gestures could be decoded from brain activity in the same two regions. Third, observing two different gestures compared with two identical gestures activated brain areas related to motor conflict, and this activity was correlated with parietal motor activity. Together, these results show that the motor system is able to represent multiple observed actions, and as such reveal a potential mechanism by which third-party social encounters could be processed in the brain.

---

There is accumulating evidence that action observation triggers a corresponding motor representation in the observer (Rizzolatti and Sinigaglia 2010, 2016; Cracco et al. 2018). For instance, animal research has uncovered a subset of motor neurons in the premotor and parietal cortex that fire both when an action is executed and when the same action is observed (Gallese et al. 1996; Kilner and Lemon 2013). This is supported by human research, in which the role of motor processes in action observation has been demonstrated across a wide range of techniques, including fMRI (Gazzola and Keysers 2009; Molenberghs et al. 2012), TMS (Fadiga et al. 1995; Naish et al. 2014), M/EEG (Arnstein et al. 2011; Fox et al. 2016), and intracranial recording (Mukamel et al. 2010; Babiloni et al. 2016).

Together, this research indicates that we represent observed actions through motor simulation in shared neural circuits (Keysers and Gazzola 2006). This, in turn, has been argued to facilitate action perception (Rizzolatti and Sinigaglia 2010, 2016). In support, studies have shown that disrupting sensorimotor brain areas impairs our ability to recognize and predict others’ actions (Avenanti et al. 2013; Urgesi et al. 2014). In social life, however, we have to represent not only the actions but also the interactions of others (Quadflieg and Koldewyn 2017; Quadflieg and Penton-Voak 2017). Recently, it was proposed that third-party interactions could be represented by simulating the actions of the different interactors in the motor system (Quadflieg and Koldewyn 2017; Quadflieg and Penton-Voak 2017). Yet, as research has mainly focused on situations with one agent, the underlying assumption that multiple observed actions can be processed in the motor system remains to be tested.

Pointing in this direction, preliminary evidence suggests that motor activation is stronger when observing an interaction between two persons than when observing a single person (Iacoboni et al. 2004; Bucchioni et al. 2013; Aihara et al. 2015) or two independently acting persons (Centelles et al. 2011; Georgescu et al. 2014). However, a non-specific increase in motor activation does not necessarily indicate that observers simulated multiple actions. Instead, to reach this conclusion, it has to be shown that multiple motor representations were activated. Therefore, in previous work, we have measured automatic imitation (Cracco et al. 2015; Cracco and Brass 2018a, 2018b) and corticospinal excitability (Cracco et al. 2016) while participants observed either a single action or two identical actions. This revealed action-specific increases in both measures when two identical actions were observed, suggesting that both actions triggered the same motor representation. Nevertheless, two identical actions might still be represented as a single action in the motor system. Therefore, a crucial question is whether it is also possible to simulate two different observed actions. Indeed, when watching two interacting individuals, it is rarely the case that their actions can be reduced to a single action.

To address this question, the current fMRI study measured sensorimotor brain activity while participants watched two right hands performing one out of three sign language gestures (see Figure 1a and Supplementary Videos). We focus on three hypotheses. First, we test whether motor activation is stronger when two hands perform two different actions (2H DIFF) than when one hand performs a single action (1H). Second, we use multivariate analyses to investigate whether both 2H DIFF gestures can be decoded from brain activation in the motor system. Finally, we explore the consequences of mirroring two different gestures. In particular, we investigate whether this leads to motor conflict, based on the fact that it is impossible to simultaneously execute two gestures with one hand. We predict stronger activation in brain areas associated with motor conflict when observing two different actions (2H DIFF) than when observing two identical actions (2H SAME). Specifically, we expect this contrast to activate the anterior cingulate cortex (ACC), which is at the core of the conflict monitoring network (Botvinick et al. 2001, 2004; Braver et al. 2001; Ridderinkhof et al. 2004).

**Figure 1.**
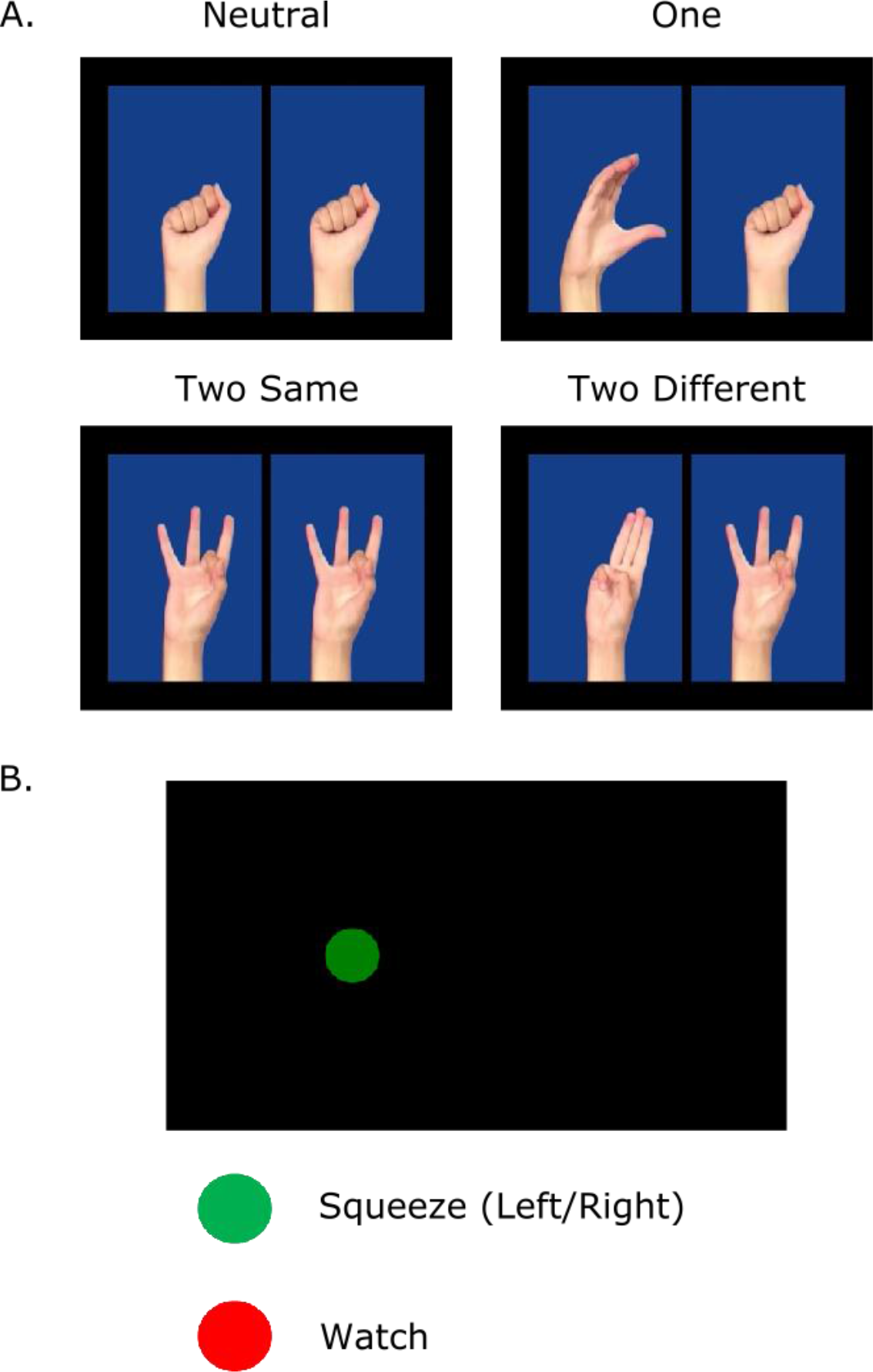
Examples of the stimuli used (a) in the action observation task and (b) in the action execution task. In the action observation task, participants watched short videos of two hands repeatedly performing one out of three possible gestures. The task of participants was to detect a glitch in the video appearing randomly in one out of seven trials. In the action execution task, participants had to either squeeze a bubble wrap ball or look at the screen depending on whether the color of the circle was, respectively, green or red. In case of a green circle, participants had to squeeze the ball each time the circle decreased in size with the hand that corresponded to the location of the circle on the screen.

## Methods

### Participants

Thirty healthy volunteers participated in the experiment for 30 euro (17 female, *M*_age_ = 22.67, *SD*_age_ = 2.48, *range*_age_ = 18 - 28), but one participant was excluded due to excessive head movement. This resulted in a sample of twenty-nine participants (17 female, *M*_age_ = 22.76, *SD*_age_ = 2.47, *range*_age_ = 18 - 28). However, the action execution data of three participants could not be used. For one participant, a technical error caused the randomization to be lost. Furthermore, for two participants, there was excessive head movement between the action observation and action execution phase, which led to missing data in the occipital lobe as a consequence of realignment. In other words, the action observation analyses were conducted on 29 participants and the action execution analyses on 26 participants. All participants were right-handed according to the Edinburgh Handedness Inventory (Oldfield 1971), did not speak sign language, had no history of neurological or psychiatric disorder, and gave written informed consent. The study was approved by the Medical Ethic Review Board of the Ghent University Hospital.

### Task and Procedure

#### Experimental Design

The experiment was structured in three phases. The first phase was a familiarization phase, which took place outside the scanner. In this phase, participants were acquainted with the two experimental tasks (i.e., the action observation and action execution tasks). Furthermore, to ensure that participants were able to execute the three gestures used in the action observation task, the familiarization phase included an imitation task in which each gesture was presented 10 times. Participants had to imitate the gesture with their right hand and then press the space bar to continue. Performance on this task was monitored by the experimenter and mistakes were corrected. Following the familiarization phase, participants were put into the scanner to complete the second and third phase of the experiment, respectively. In the second phase, participants performed the action observation task. In the third phase, they performed the action execution task.

#### Action Observation Task

In the action observation phase, participants watched short videos (5 s) of two right hands repeatedly (4 times) performing one out of three sign language gestures (Fig. 1a and Supplementary Videos). The gestures were chosen on the basis of a pilot study in which 40 participants used a 5-point Likert scale to rate eight sign language gestures on whether the gesture was familiar, had been seen before, was clearly visible, was difficult to execute, and had a meaning. For each gesture, participants were furthermore asked to guess its meaning. Based on the results, we then chose three gestures that were clearly visible, not too difficult to execute, and with unknown meaning. The videos were presented as a sequence of 28 frames. The first frame showed the hands in rest position and was presented for 300 ms. Next, all 28 frames were presented for 33 ms each. Finally, the last frame remained on the screen for another 300 ms before the 28 frames were presented again for 33 ms each. This was repeated for four cycles so that each video had a total duration of 5196 ms (300 ms + 4 x 28 x 33 ms + 4 x 300 ms).

To ensure that attention was maintained throughout the entire duration of the experiment, participants had to detect a glitch appearing randomly in one seventh of the trials at a random moment in the video on the left hand, on the right hand, or on both hands. The glitches were inserted by replacing 1 of the 28 frames with a blue frame. This resulted in a brief flicker (33 ms) that was easily missed if attention was not focused on both hands (Video S4). After each glitch trial, two questions were presented on the screen. The first question required participants to indicate the gesture(s) on which the glitch had appeared. That is, a series of four pictures was presented showing the neutral hand followed by the three gestures. For each picture, participants had to indicate whether a glitch had appeared on the presented gesture or not. The second question asked participants whether the glitch had appeared on one hand or on both hands. Accuracy was 75% on the first question and 85% on the second question, indicating that the task was challenging but not too difficult. All participants scored above chance level on both questions. Trials with a glitch were not included in the analyses.

The action observation task comprised two runs of 126 trials each. Trials were presented randomly with the restriction that all 18 gesture combinations occurred equally often in both runs. The following gesture combinations were included in the experiment: nG1/nG1, nG2/nG2, nG3/nG3, nG1/G1, G1/nG1, nG2/G2, G2/nG2, nG3/G3, G3/nG3, G1/G1, G2/G2, G3/G3, G1/G2, G2/G1, G1/G3, G3/G1, G2/G3, G3/G2, with G1, G2, and G3 representing the three gestures and nG1, nG2, and nG3 representing the corresponding neutral hand. For example, G2/G3 means that gesture 2 was presented left and gesture 3 was presented right. The 18 gesture combinations were combined to form four conditions. In the Neutral condition (N), neither hand performed a gesture. In the One Hand condition (1H), one hand performed a gesture. In the Two Hands Same condition (2H SAME), both hands performed the same gesture. Finally, in the Two Hands Different condition (2H DIFF), the two hands each performed a different gesture. Note that the N and 2H SAME conditions contained fewer gesture combinations and therefore fewer trials than the 1H and 2H DIFF conditions. However, they still included 36 trials each, thereby providing us with sufficient data to reliably estimate the signal in all conditions. All trials were separated by a black screen presented for a jittered duration drawn from a pseudo-logarithmic distribution with 50% short durations (200, 800, 1400, or 2000 ms), 33.3% intermediate durations (2600, 3200, 3800, or 4400 ms), and 16.7% long durations (5000, 5600, 6200, or 6800 ms).

#### Action Execution Task

Immediately after the action observation task, participants performed the action execution task. In this task, a green or red circle was presented on the left or right side of the screen. The circle then decreased in size every second until it disappeared after four seconds. During the action execution task, participants wore gloves with a bubble wrap ball attached to the palm. When a green circle was presented, participants had to squeeze the ball each time the circle decreased in size using the hand that corresponded to the location of the circle on the screen. For example, when a green circle appeared on the left side, participants had to squeeze the ball using their left hand. In contrast, when a red circle appeared, participants simply had to watch the screen. The action execution task consisted of 60 trials that were randomly subdivided into 20 squeeze left trials, 20 squeeze right trials, and 20 watch trials. Trials were separated by a black screen presented for a jittered duration of 4000, 5000, 6000, 7000, or 8000 ms. The action execution task was based on the task used by Arnstein et al. (2011). Importantly, these authors found that a simple squeeze task like the one used here activated a network very similar to the network activated by a more complex object manipulation task.

### fMRI Parameters

MRI images were acquired with a 3T Siemens Trio scanner using a 32-channel radiofrequency head coil. Participants entered the scanner head first and supine. The scanning procedure started with an anatomical scan in which 176 high-resolution anatomical images were acquired using a T1-weighted 3D MPRAGE sequence [repetition time (TR) = 2250 ms, echo time (TE) = 4.18 ms, image matrix = 256 x 256, field of view (FOV) = 256 x 256 mm, flip angle = 9°, voxel size = 1.00 x 1.00 x 1.00 mm]. This was followed by two action observation runs and one action execution run in which whole-brain functional images were obtained. These functional images were acquired using a T2*-weighted echo planar imaging (EPI) sequence, sensitive to BOLD contrast (TR = 2000 ms, TE = 28 ms, image matrix = 64 x 64, FOV = 224 mm, flip angle = 80°, distance factor = 17%, voxel size 3.5 x 3.5 x 3.0 mm, 34 axial slices).

### Analyses

#### fMRI Preprocessing

All fMRI data was analyzed using SPM8 (Wellcome Department of Imaging Neuroscience, UCL, London, U.K.; www.fil.ion.ucl.ac.uk/spm). To account for T1 relaxation, the first four scans of all runs were dummy scans. The three runs were preprocessed together according to the following steps. First, the functional images were spatially realigned using a rigid body transformation. Second, the realigned images were slice-time corrected with respect to the middle acquired slice. Third, the structural image of each subject was co-registered with its mean functional image. Fourth, the anatomical images were segmented according to the SPM8 tissue probability maps and the resulting parameters were used to normalize the functional images to MNI space. Finally, the images were resampled into 3 mm^3^ voxels and spatially smoothed with an 8 mm Gaussian kernel (full-width at half maximum). However, for the representational similarity analyses, we used the unsmoothed data instead.

#### Univariate Analyses

The data was filtered using a 128 Hz high-pass filter. First level analyses were conducted by fitting a general linear model in SPM8. The action observation model consisted of 10 regressors per run, namely one regressor for each condition (N, 1H, 2H SAME, and 2H DIFF) and six regressors representing the realignment parameters. The signal was modeled over the entire duration of the videos and was convolved with the canonical hemodynamic response function (HRF). The action execution model also consisted of 10 regressors, with one regressor for each condition (Watch Left, Watch Right, Squeeze Left, Squeeze Right) and six regressors representing the realignment parameters. The entire duration of the trial, from the moment the circle appeared until the moment it disappeared, was modeled and convolved with the canonical HRF.

The second level analyses of both tasks were performed using a within-subject one-way ANOVA with equal variances. Unless otherwise specified, all results are thresholded using a *p* < .05 FWE-corrected threshold at the peak level. Whole brain univariate analyses were used to localize action execution activation, action observation activation, shared voxel (sVx) activation, and motor conflict activation. To localize action execution activation (Squeeze > No Squeeze), we calculated the contrast [Squeeze Left + Squeeze Right] > [Watch Left + Watch Right]. To localize action observation activation (Observation > Neutral), we calculated the contrast 0.33 x [1H + 2H SAME + 2H DIFF] > N. The intersection of these two activation maps was used to localize sVx activation, which served as a proxy for the human mirror neuron system (Gazzola and Keysers 2009). Finally, to localize motor conflict activation, we calculated the contrast 2H DIFF > 2H SAME.

#### Region of Interest Analyses

To test whether brain activity was stronger in the 2H DIFF condition than in the 1H condition, we constructed three regions of interest (ROIs) from brain activity in the action observation task. To ensure that the ROIs were not biased towards the 2H conditions, we defined them using the Nichols et al. (2005) conjunction contrast [1H > N] ∩ [2H SAME > N] ∩ [2H DIFF > N] (Figure S1). Moreover, to secure statistical independence, we used a leave-one-out cross-validation procedure in which the ROIs for each participant were calculated using the data of all participants except that one participant (Esterman et al. 2010). That is, for each participant, we performed the specified conjunction analysis on the data of all the other participants. We then defined the ROIs by constructing 5 mm spheres around the peak coordinates of the left and right visual area 5 (V5), the left and right parietal cortex (PAR), and the left premotor cortex (PMC). The right PMC was not consistently activated. Therefore, to obtain a bilateral PMC ROI, we mirrored the left PMC sphere onto the right hemisphere. Importantly, the activation pattern was highly similar in the left and right PMC (Figures S2-S3).

The V5 peak coordinates were determined using the Squeeze > No Squeeze action execution activation as an exclusive mask (*p* < .001, uncorrected). Conversely, the PAR and PMC peak coordinates were determined using the Squeeze > No Squeeze action execution activation as an inclusive mask (*p* < .001, uncorrected). The use of an uncorrected *p* < .001 threshold for the masks was based on previous work (Arnstein et al. 2011). Moreover, we intersected the obtained spheres with a brain mask to ensure that the ROIs did not extend beyond the brain. The peak coordinates used to construct the ROIs are reported separately for each participant in Table S1. To perform the univariate ROI analyses, we extracted the beta values using the MARSBAR package in SPM8 (Brett et al., 2002), added the intercept to each value, and then used the resulting values to calculate the percent signal change in the 1H and 2H DIFF conditions with respect to the N condition as 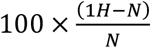 and 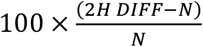.

#### Representational Similarity Analysis

To investigate whether both 2H DIFF gestures could be decoded from brain activity in the three ROIs, we conducted a representational similarity analysis (Kriegeskorte et al. 2008), which was performed on the unsmoothed data. First level analyses were conducted by fitting a general linear model in SPM8. The model consisted of 24 regressors per run, namely one regressor for each of the 18 gesture combinations and six regressors for the realignment parameters. The signal was modeled over the entire duration of the videos and was convolved with the canonical HRF. Next, for each voxel in each ROI, we extracted the beta values corresponding to the 18 gesture combinations and then computed pairwise correlations across voxels to obtain an 18 x 18 correlation matrix per run per ROI. In these matrices, each cell compares the activation pattern of two gesture combinations (e.g., G1/G2 and G3/G3). To account for the non-normal distribution of the correlation coefficients, we first transformed the correlations into Fisher *z*-scores before using them in the analyses.

To decode the 2H DIFF gestures from brain activity in the three ROIs, we correlated the activation pattern in the 2H DIFF condition with the activation patterns in the 1H and 2H SAME conditions. This allowed us to compare two correlations with gesture overlap and one correlation without gesture overlap (Fig 4a). That is, it allowed us to compare the activation pattern when observing both gestures A and B with the activation pattern when observing only gesture A (Overlap A), only gesture B (Overlap B), or only gesture C (No Overlap). Importantly, the representational similarity analyses were conducted without considering the location of the gestures on the screen, meaning that Overlap A was the mean of r[A/B, nA/A], r[A/B, A/nA], and r[A/B, A/A], Overlap B was the mean of r[A/B, nB/B], r[A/B, B/nB], and r[A/B, B/B], and No Overlap was the mean of r[A/B, nC/C], r[A/B, C/nC], and r[A/B, C/C]. The above correlations were calculated separately for the different possible gesture combinations in the 2H DIFF condition (G1/G2, G2/G1, G1/G3, G3/G1, G2/G3, and G3/G2), and were then averaged so that one score was obtained per participant per Overlap condition.

#### Psychophysiological Interaction Analysis

Finally, we conducted a psychophysiological interaction (PPI) analysis using the gPPI toolbox (McLaren et al. 2012) to investigate whether motor conflict activation depended on sVx activation. The ACC, which was identified using the contrast 2H DIFF > 2H SAME in the action observation task, was used as seed region. PPI regressors were calculated at the first level for all four conditions (N, 1H, 2H SAME, and 2H DIFF) and were then analyzed using a within-subject one-way ANOVA with equal variances at the second level. Because we were only interested in connections with the action observation network, the PPI analysis was restricted to voxels that were significant in the action observation task (Observation > Neutral) at an uncorrected *p* < .001 threshold. Within this volume, the PPI results were thresholded at FWE-corrected *p* < .05 peak threshold.

## Results

### Shared Voxel Localization

Shared voxel activation was determined by measuring the overlap in brain activation between action observation and action execution. First, to localize action execution activation, we calculated the contrast Squeeze > No Squeeze. As shown in Figure 2a, this revealed widespread activation in the sensorimotor system, consistent with previous research using this task (Arnstein et al. 2011). Next, to localize action observation activation, we compared activation in the N condition with activation in the 1H, 2H SAME, and 2H DIFF conditions using the contrast Observation > Neutral. As expected, this yielded bilateral activation around V5, and in the inferior parietal cortex, the superior parietal cortex, the postcentral gyrus, and the superior temporal gyrus, as well as lateralized activation in the left dorsal premotor cortex (Fig 2b). Finally, to localize sVx, we determined the overlap between both tasks. In line with meta-analyses on action observation (Caspers et al. 2010; Molenberghs et al. 2012), this revealed sVx activation in the inferior parietal cortex, the superior parietal cortex, the postcentral gyrus, the superior temporal gyrus, and the dorsal premotor cortex, but not in the visual cortex (Fig 2c).

**Figure 2.**
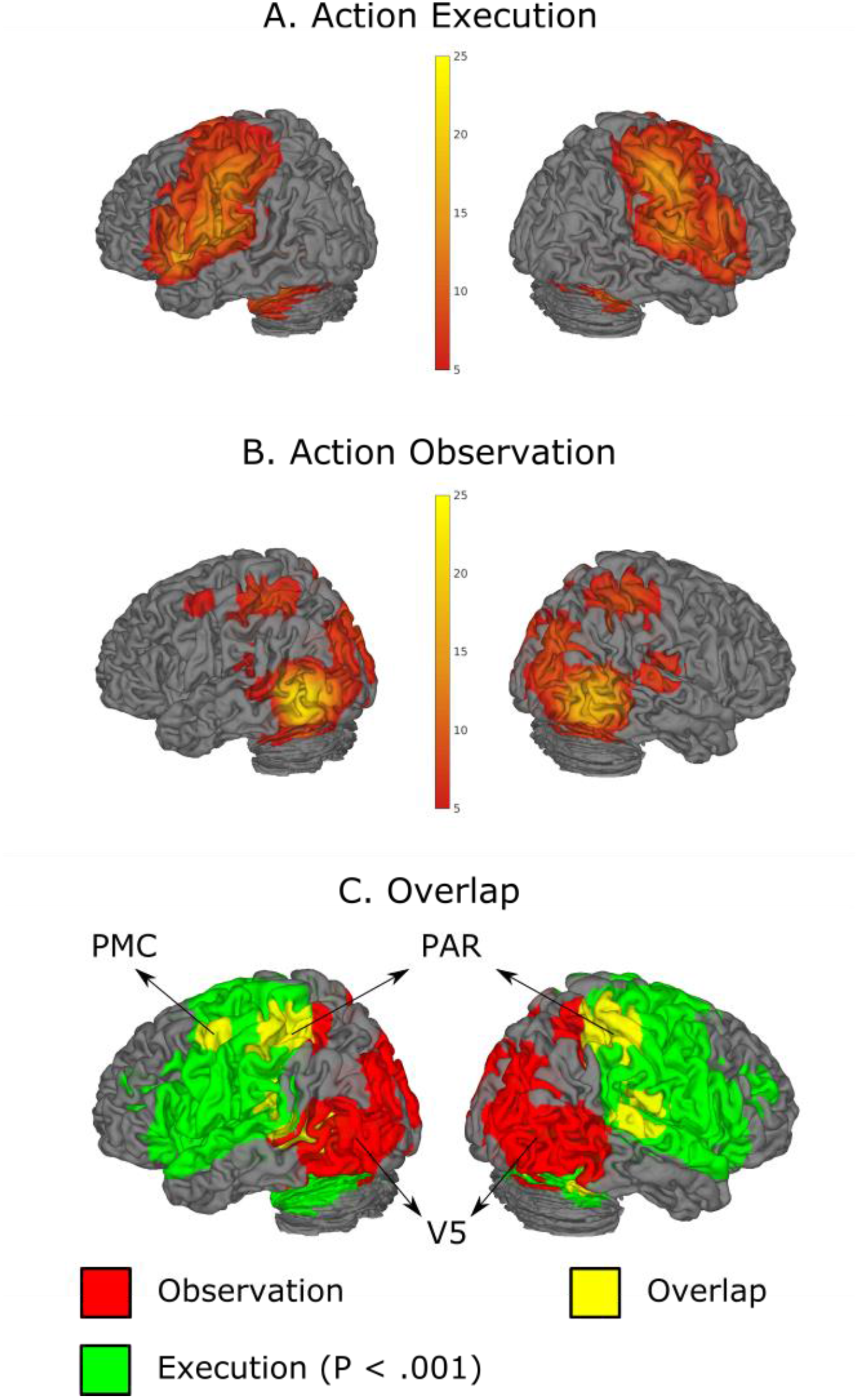
The top panel shows brain activation in the action observation task (Observation > Neutral). The middle panel shows brain activation in the action execution task (Squeeze > No Squeeze). The bottom panel shows the overlap between both tasks. In line with previous work, the action execution data was thresholded at *p* < .001 (uncorrected) to determine shared voxels (Arnstein et al. 2011). All other images were thresholded using a *p* < .05 FWE-corrected threshold

### Representing Two Observed Actions in the Motor System

#### Is There More Activation?

To test whether motor activation was stronger when observing two different actions compared with a single action, we looked at brain activity in three core regions of the action observation network, namely one visual region (V5) and two sensorimotor regions (PAR, and PMC). That is, we conducted a region x condition repeated measures MANOVA to compare the percent signal change in the 1H condition with the percent signal change in the 2H DIFF condition, both relative to the N condition, averaged across the left and right hemisphere (but see Figures S1 and S2 for the hemisphere specific results). As predicted, this revealed a main effect of condition, *F*(1, 28) = 79.15, *p* < .001, indicating stronger activity in the 2H DIFF condition than in the 1H condition. The region x condition interaction was significant as well, *F*(2, 27) = 86.06, *p* < .001 (Fig 3). This showed that the condition effect was stronger in V5 than in both PAR, *t*(28) = 11.35, *p* < .001, and PMC, *t*(28) = 13.34, *p* < .001. However, in addition to V5, *F*(1, 28) = 155.60, *p* < .001, there was also a significant effect of condition in PAR, *F*(1, 28) = 16.40, *p* < .001, and PMC, *F*(1, 28) = 5.56, *p* = .026. In other words, these results indicate that brain activity in visual as well as motor areas was stronger when two different gestures were observed compared with when a single gesture was observed. As such, they support the hypothesis that both 2H DIFF gestures were processed in the motor system.

**Figure 3.**
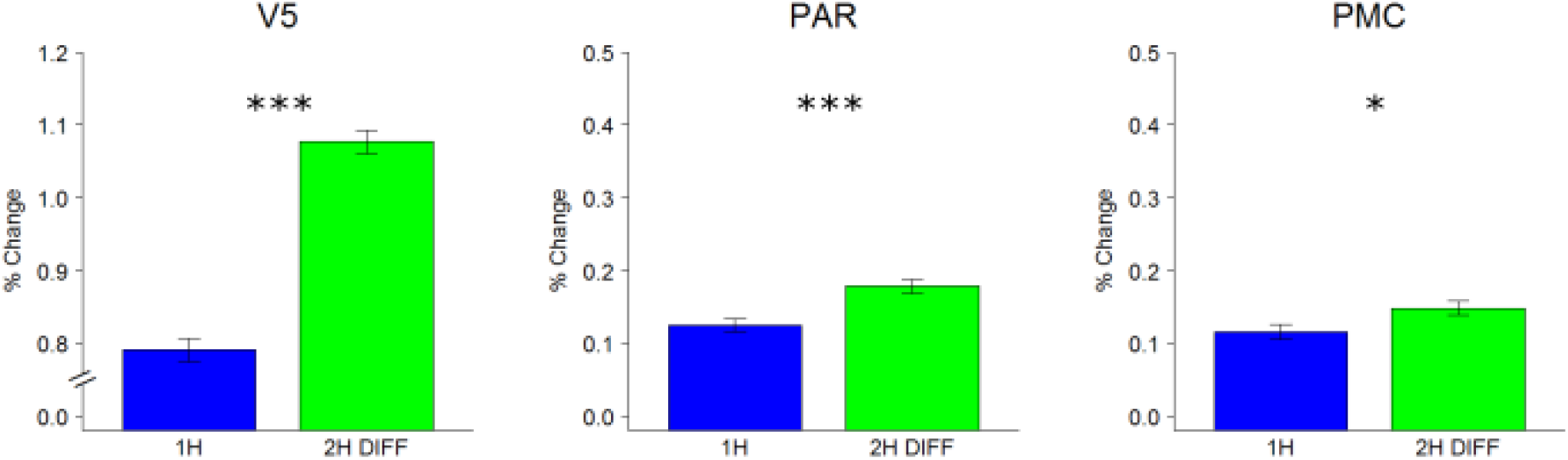
Results of the ROI analyses testing whether activation was stronger in the 2H DIFF condition than in the 1H condition. The y-axis shows the % signal change with respect to the N condition. Details on the % signal change calculation are provided in the methods. Post-hoc two-tailed *t* tests comparing 1H with 2H DIFF are displayed. Error bars are standard errors of the mean (SEMs) corrected for within-subject designs (Morey 2008). * p < .05; ** p < .01; *** p < .001.

#### Can We Decode Both Actions?

To further inform whether participants represented both 2H DIFF gestures in their motor system, we then tried to decode these gestures from brain activity in the three ROIs using representational similarity analysis, which is a multivariate technique to measure the similarity between the representations of two stimuli by correlating their activation patterns (Kriegeskorte et al. 2008). Applied to the current study, we correlated the activation pattern in the 2H DIFF condition with the activation pattern in the 1H and 2H SAME conditions because this allowed us to compare two correlations with gesture overlap and one correlation without gesture overlap (Figure 4a). More specifically, it allowed us to compare the activation pattern when observing both gesture A and gesture B with the activation pattern when observing only gesture A (Overlap A), only gesture B (Overlap B), or only gesture C (No Overlap).

**Figure 4.**
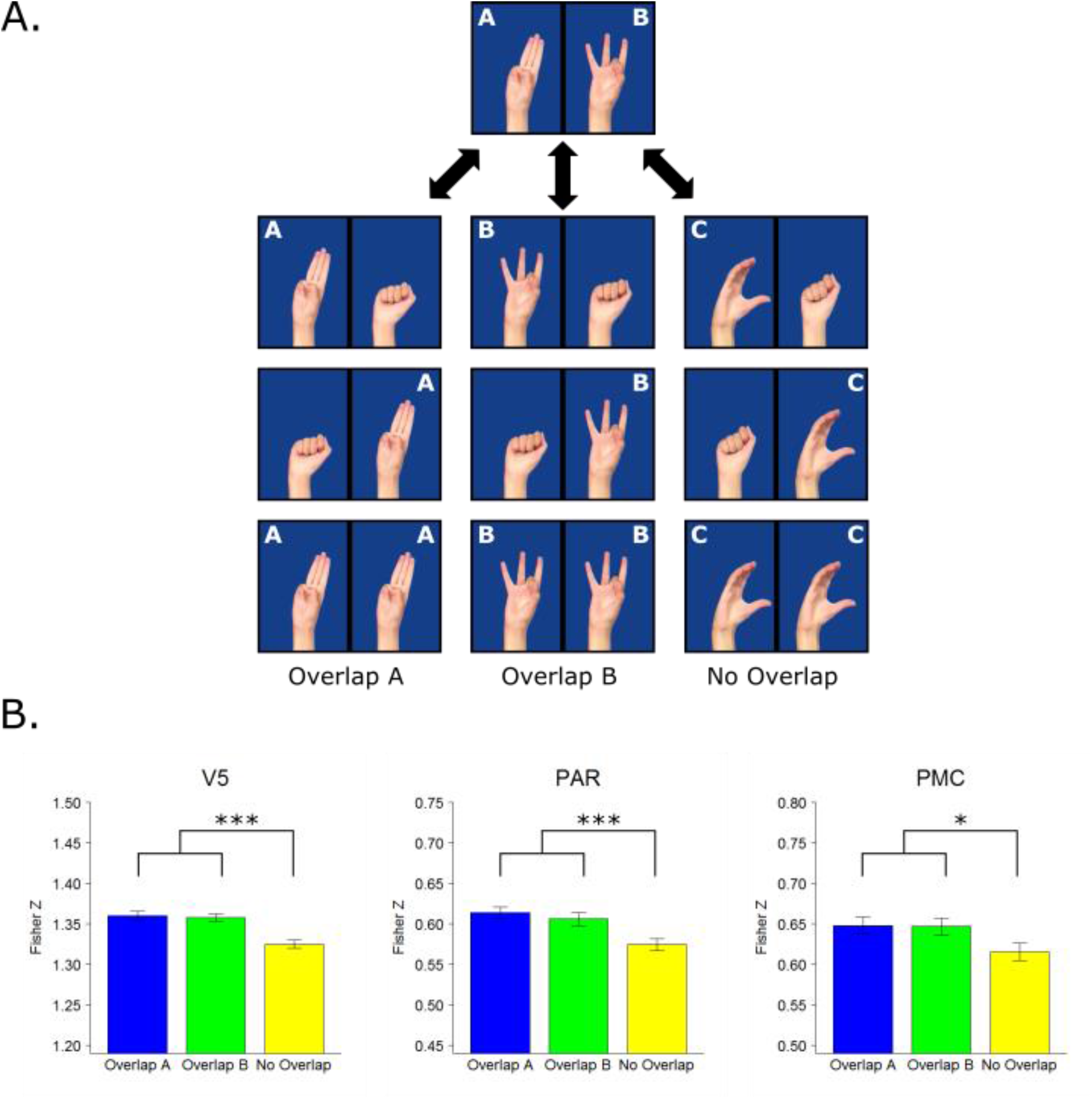
Results of the representational similarity analyses testing whether the two gestures observed in the 2H DIFF condition could be simultaneously decoded from brain activation in the three ROIs. Panel A is a visual representation of the analysis. In the Overlap A condition, gesture A overlaps. In the Overlap B condition, gesture B overlaps. In the No Overlap condition, neither of the two gestures overlaps. The analysis disregards the location of the gestures on the screen. Panel B shows the Fisher *Z*-transformed correlation coefficients in each of the three conditions separately for each ROI. Post-hoc two-tailed *t* tests comparing the average of Overlap A and B with No Overlap are displayed. The difference between Overlap A and Overlap B was never significant. Error bars are SEMs corrected for within-subject designs (Morey 2008). * p < .05; ** p < .01; *** p < .001.

If participants represented both gestures in the 2H DIFF condition, the correlations should be stronger when there is gesture overlap than when there is no gesture overlap. To evaluate this hypothesis, we conducted a region x overlap repeated measures MANOVA on the Fisher *Z*-transformed correlations in the three ROIs (Fig. 4b). As predicted, this revealed a main effect of overlap, *F*(2, 27) = 11.63, *p* < .001, with stronger spatial correlations in the Overlap A condition than in the No Overlap condition, *t*(28) = 4.70, *p* < .001, stronger correlations in the Overlap B condition than in the No Overlap condition, *t*(28) = 3.96, *p* < .001, but no difference between the Overlap A and Overlap B conditions, *t*(28) = 0.53, *p* = .601 (Fig 4b). There was no region x overlap interaction, *F*(4, 25) = 0.07, *p* = .991. Taken together, these results thus indicate that the actions of both hands could be decoded from brain activity in visual as well as motor areas even when the hands performed different actions.

### Does Representing Multiple Observed Actions Produce Motor Conflict?

Finally, we looked at the consequences of mirroring multiple observed actions. That is, because the 2H DIFF gestures are mutually exclusive in terms of motor execution, mirroring them should lead to motor conflict (Botvinick et al. 2001, 2004) and therefore to ACC activation (Botvinick et al. 2001, 2004; Braver et al. 2001; Ridderinkhof et al. 2004). To test this hypothesis, we compared brain activity in the 2H DIFF condition with brain activity in the 2H SAME condition (2H DIFF > 2H SAME). As expected, this activated the ACC. In addition, it also activated the right anterior insula, which is known to co-activate with the ACC (Ullsperger et al. 2014), and the action observation network (Fig 5; Table 1). Next, to evaluate whether this corresponds to what is typically found in motor conflict research, we performed a Neurosynth meta-analysis on the search string (*Response*\*/*Motor*) & (*Conflict*\*/ *Compet*\*). That is, we computed reverse inference maps to investigate whether the ACC and AI were activated more consistently in studies mentioning motor conflict than in studies not mentioning motor conflict (Yarkoni et al. 2011). As shown in Fig 5, this was indeed the case, and an overlap analysis revealed substantial overlap with the pattern of activation obtained in the current study.

**Figure 5.**
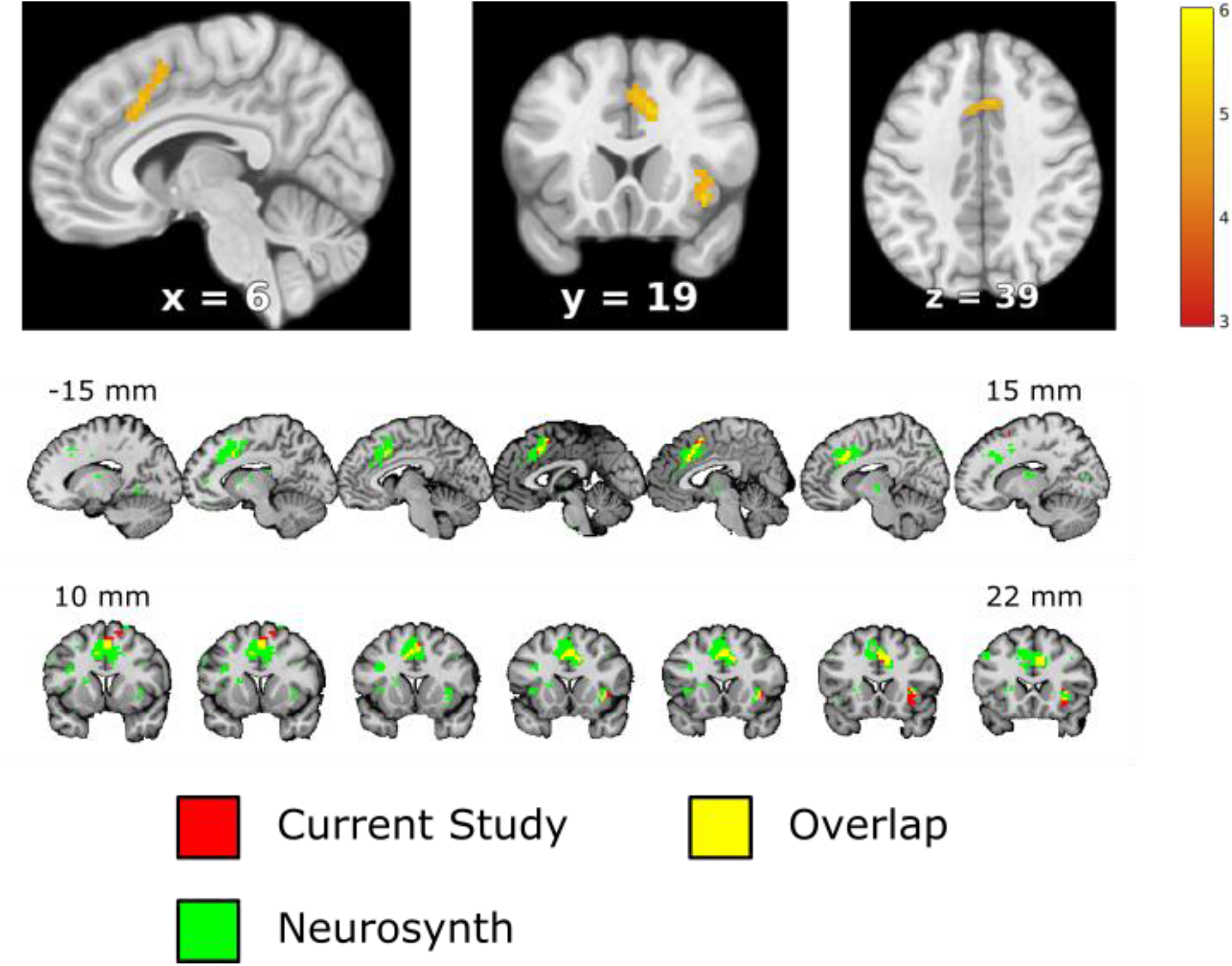
The top panel shows the univariate results of the 2H DIFF > 2H SAME contrast in the action observation task. The bottom panel shows the overlap between the results of the current study and the results of a Neurosynth meta-analysis on motor conflict. Brain activation from the current study is thresholded at *p* < .05 using FWE correction. Brain activation from Neurosynth is thresholded at *p* < .01 using FDR correction.

**Table 1.** Peak MNI Coordinates of the 2H DIFF > 2H SAME contrast.

| Brain area | Peak coordinates | T-score | Cluster size |
| --- | --- | --- | --- |
| ACC, Pre-SMA | 12, 20, 34 | 5.42 | 85 |
|  | 3, 11, 52 | 5.07 |  |
|  | 3, 17, 43 | 5.05 |  |
| Right AI | 36, 20, -8 | 5.37 | 28 |
|  | 42, 17, -2 | 5.01 |  |
|  | 33, 23, 4 | 4.95 |  |
| Right IPG | 30, -49, 46 | 5.07 | 7 |
| Left dPMC | -33, -4, 52 | 5.06 | 8 |
| Right MTG | 45, -70, 13 | 5.00 | 6 |
| Right SFG | 12, 11, 64 | 4.89 | 4 |
| Right dPMC | 36, -4, 55 | 4.82 | 8 |
|  | 42, 2, 58 | 4.77 |  |

*Note.* ACC = Anterior Cingulate Cortex, pre-SMA = Pre-supplementary Motor Area, AI = Anterior Insula, IPG = Inferior Parietal Gyrus, dPMC = dorsal premotor cortex, MTG = middle temporal gyrus, SFG = superior frontal gyrus. All results are FWE-corrected at $p < .05$ .

Crucially, however, if the ACC activity was caused by motor conflict, then it should depend on activity in the motor system. That is, according to the response conflict model of cognitive control, the ACC detects and signals the presence of motor conflict (Botvinick et al. 2001, 2004). In particular, this model argues that response conflict can be seen as the product of the activation in two simultaneously active response nodes. As a consequence, if there is only one active response node, there is no response conflict. In contrast, if there are two active response nodes, response conflict depends on the activation of both nodes. In the current study, two identical observed gestures should trigger the same “response”, whereas two different observed gestures should trigger two different “responses”. Therefore, response conflict is expected in the 2H DIFF but not in the 2H SAME condition, and response conflict in the 2H DIFF condition should depend on the strength of activation in the motor system.

To test this hypothesis, we calculated the PPI of the 2H DIFF > 2H SAME contrast using the ACC as seed region and the action observation activation (Observation > Neutral) as an inclusive small volume corrected mask (Fig 6). This revealed a condition-dependent association with activity in the inferior parietal cortex, and this activity completely overlapped with the activity in the action execution task (Squeeze > No Squeeze). Thus, the PPI analysis showed that activity in parietal sVx was correlated more strongly with ACC activity in the 2H DIFF condition than in the 2H SAME condition. In line with the motor conflict hypothesis, this suggests that ACC activity signaled the presence of motor conflict produced by simultaneously representing two mutually exclusive observed gestures in the motor system.

**Figure 6.**
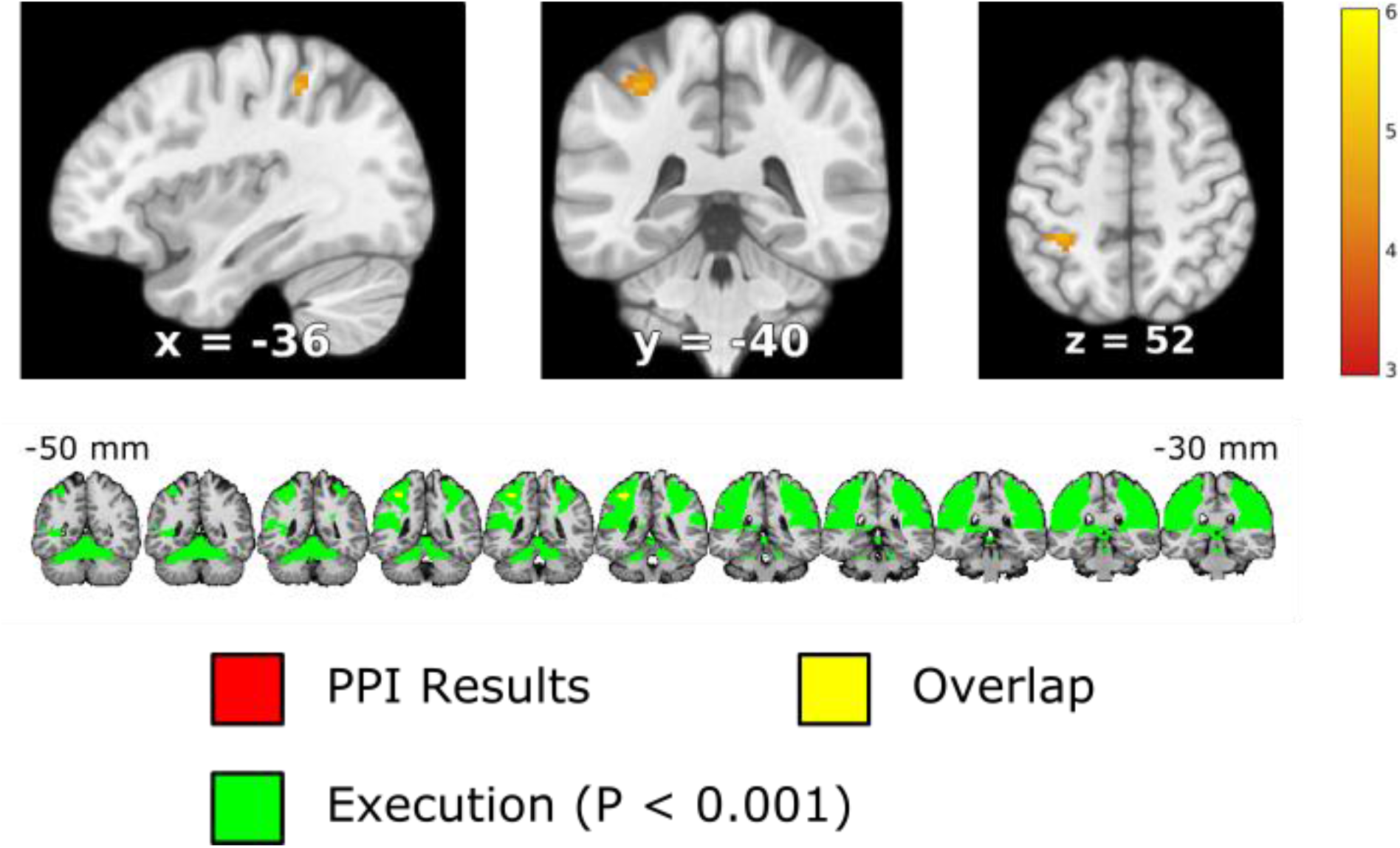
The top panel shows the PPI results of the 2H DIFF > 2H SAME contrast in the action observation task. The displayed coordinates are the peak coordinates of the PPI analysis (*t* = 4.99, *k* = 20). Note that the PPI results are masked with the activation of the action observation task (Observation > Neutral). The bottom panel shows the overlap between the activation of the PPI analysis and the activation of the action execution task (Squeeze > No Squeeze). There was no PPI activation that was not captured by the action execution activation. Unless otherwise specified, brain activation is thresholded with a small-volume FWE-corrected *p* < .05 threshold. In line with previous work, action execution data was thresholded at *p* < .001 to determine activation overlap (Arnstein et al. 2011).

## Discussion

The mirror mechanism is a fundamental neural mechanism that translates observed actions into motor programs (Rizzolatti and Sinigaglia 2010, 2016). It has been argued that this mechanism supports action representation through motor simulation (Avenanti et al. 2013; Urgesi et al. 2014). However, in contrast to action simulation, little is known about interaction simulation. A necessary condition for interaction simulation is that multiple observed actions can be represented in the motor system (Quadflieg and Koldewyn 2017; Quadflieg and Penton-Voak 2017). In the current study, we investigated this condition by asking participants to watch two right hands performing sign language gestures. Three key results were obtained. First, the motor system was activated more strongly when two different gestures were observed compared with when a single gesture was observed. Second, both individual gestures could be decoded from activity in the motor system. Third, observing two different gestures relative to observing two identical gestures produced motor conflict related activity in the ACC, and this activity was correlated more strongly with parietal motor activity in the former condition than in the latter condition. Together, these results indicate that the motor system is able to simultaneously represent multiple observed actions, even when they cannot be simultaneously executed. Instead, when the mirrored actions violate motor constraints, this is signaled in the form of motor conflict.

Nevertheless, an alternative explanation is that participants did not represent both gestures simultaneously, but instead randomly represented one gesture on each trial. In this case, the activation pattern over trials would combine both gestures, likewise making it possible to decode both gestures from the average brain activation. However, a random sampling mechanism seems unlikely considering that our attentional task required participants to simultaneously monitor both hands. Indeed, performance on this task (85%) was well above chance (50%). In the same vein, random sampling is inconsistent with recent evidence that non-attended observed actions modulate automatic imitation even when another observed action is being attended or being imitated (Cracco and Brass 2018a). Finally, random sampling is difficult to reconcile with the other two main results, namely that observing two different gestures produced stronger motor responses and led to motor conflict. That is, if only one gesture was represented on each trial, then motor activation should not be stronger when two gestures were observed, nor should there be any motor conflict. Indeed, motor conflict is widely held to arise from the simultaneous processing of two or more actions (Botvinick et al. 2001). As a result, it should only be present if both actions were represented in parallel.

Still, one could argue that the interpretation of ACC activation as motor conflict was a reverse inference and therefore invalid. However, it should be stressed that this interpretation was not a post-hoc explanation of unexpected brain activation but rather an a-priori hypothesis restricted by theoretical as well as task constraints. Speaking more broadly, the validity of reverse inferences critically depends on the likelihood that the proposed process, as opposed to competing processes, was engaged in the experimental task (Poldrack 2006; Hutzler 2014). Looking at the literature, two plausible alternative processes can be identified. First, it could be argued that the ACC was activated due to differences in the frequency with which certain stimuli were presented, consistent with evidence that this region codes expectancy violation (Desmet et al. 2014; Fouragnan et al. 2018). However, this should have produced the opposite pattern of results, considering that 2H SAME trials (i.e., G1/G1, G2/G2, and G3/G3) occurred less often than 2H DIFF trials (i.e., G1/G2, G2/G1, G1/G3, G3/G1, G2/G3, and G3/G2).

Second, it could be argued that ACC activity was driven by stimulus conflict rather than motor conflict. As stimulus conflict occurs already at the visual level (Verbruggen et al. 2006), it does not require both observed gestures to be represented in the motor system. Importantly, however, an explanation in terms of stimulus conflict is inconsistent with previous research on the neural substrates of conflict processing. In particular, this work has demonstrated that the ACC is sensitive to motor conflict, but not to stimulus conflict (Milham et al. 2001; van Veen et al. 2001; Liu et al. 2004; Liston et al. 2006; Wendelken et al. 2009; Mayer et al. 2012). Furthermore, stimulus conflict cannot easily explain why ACC activation was correlated with parietal sVx activation but not with purely visual activation. That is, if stimulus conflict was coded in the ACC, it should co-activate with the visual cortex (Botvinick et al. 2001). More generally, potential alternative explanations are only feasible if they are also able to explain why ACC activation was correlated with motor activation, and why this correlation was stronger when the hands performed two different gestures than when they performed two identical gestures. Since this pattern was derived directly from conflict monitoring theories (Botvinick et al. 2001, 2004), it strongly favors the motor conflict hypothesis (Poldrack 2006; Hutzler 2014).

In sum, the current study shows that at least two observed actions can be represented at the same time in the motor system. This critically extends research on mirror processes from action representation (Rizzolatti and Sinigaglia 2010, 2016) to interaction representation (Quadflieg and Koldewyn 2017; Quadflieg and Penton-Voak 2017). That is, in contrast to what might intuitively be expected, it shows that motor simulation does not break down when observing multiple actions. Instead, in those situations, the different actions are simulated together. The ability to mirror multiple observed actions fulfills the *condicio sine qua non* of interaction simulation. As such, the present work opens up the possibility that mirror processes contribute not only to decoding the actions (Avenanti et al. 2013; Urgesi et al. 2014) but also to decoding the interactions of others (Quadflieg and Koldewyn 2017; Quadflieg and Penton-Voak 2017). However, to fully evaluate this hypothesis, it is imperative to broaden the current research from observing isolated actions to observing meaningful social interactions.

Moreover, in addition to interaction representation, the present work has important implications for joint action. That is, previous research has shown that motor simulation facilitates interpersonal coordination in tasks that require two persons to cooperate (Colling et al. 2013; Kourtis et al. 2013). For example, a recent study found that musicians’ ability to tune their own actions to those of another musician in a musical duet was impaired when the dorsal premotor cortex was disturbed (Hadley et al. 2015). However, social tasks often extend beyond the dyad (Tsai et al. 2011). For instance, musicians in a musical ensemble have to coordinate their actions not just with one but with multiple co-musicians (Volpe et al. 2016). The results of the present study suggest that this may likewise rely on motor simulation. In particular, it suggests that mirror processes could be used to simultaneously predict the action outcomes of several co-actors (Aglioti et al. 2008; Hamilton and Grafton 2008) to achieve interpersonal coordination in multi-agent settings (Colling et al. 2013; Kourtis et al. 2013).

Finally, to the best of our knowledge, the current study is the first demonstration that motor conflict is not restricted to action planning (Botvinick et al. 2001, 2004), but can occur during action observation as well. This has important implications for theories of cognitive control. For example, a prominent view is that motor conflict signals the need to increase attentional control (Botvinick et al. 2001, 2004). Thus, in this view, when observers experience motor conflict, this could trigger compensatory mechanisms aimed at increasing attention towards the observed actions, and this could then facilitate social processes such as interaction understanding (Quadflieg and Koldewyn 2017; Quadflieg and Penton-Voak 2017) or interpersonal coordination (Colling et al. 2013; Kourtis et al. 2013).

It is also interesting to note that motor conflict did not activate the temporo-parietal junction (TPJ). The TPJ is known to respond to conflict between observed and planned actions (Brass et al. 2005, 2009), and is therefore believed to distinguish self-from other-generated actions (Brass et al. 2009). As a consequence, it could be expected to distinguish between other-generated actions as well. However, self-other and other-other distinction do not necessarily rely on the same processes (Cracco and Brass 2018a). Furthermore, because no response was needed, our task did not require participants to distinguish between action representations. Thus, in this view, the ACC and TPJ may fulfill complementary roles with the ACC being involved in signaling motor conflict (Botvinick et al. 2001) and the TPJ in solving the conflict when the need for an overt action requires it (Brass et al. 2009).

To conclude, the present work demonstrates that multiple observed actions can be represented at the same time in the motor system. This has important implications for interaction representation as well as joint action, and opens up new hypotheses on the role of motor conflict in action observation.

## Supplementary Information

**Table S1.** Center MNI coordinates for the three ROIs per participant.

|  | Left V5 | Right V5 | Left PAR | Right PAR | Left PMC | Right PMC |
| --- | --- | --- | --- | --- | --- | --- |
| Subject 1 | -51, -76, -2 | 51, -70, -2 | -30, -43, 52 | 30, -40, 46 | -42, -4, 58 | 42, -4, 58 |
| Subject 2 | -51, -73, 1 | 51, -70, -2 | -33, -43, 55 | 30, -40, 49 | -42, -7, 58 | 42, -7, 58 |
| Subject 3 | -51, -76, -2 | 51, -70, -2 | -30, -43, 52 | 30, -40, 46 | -42, -4, 58 | 42, -4, 58 |
| Subject 4 | -51, -76, -2 | 51, -67, -2 | -30, -43, 52 | 30, -40, 46 | -42, -4, 58 | 42, -4, 58 |
| Subject 5 | -51, -76, -2 | 51, -70, 2 | -33, -43, 55 | 30, -43, 52 | -42, -4, 58 | 42, -4, 58 |
| Subject 6 | -51, -76, -2 | 51, -70, -2 | -30, -46, 52 | 30, -40, 46 | -42, -4, 58 | 42, -4, 58 |
| Subject 7 | -51, -76, -2 | 51, -67, -2 | -30, -43, 52 | 30, -37, 46 | -42, -4, 58 | 42, -4, 58 |
| Subject 8 | -51, -76, -2 | 51, -70, -2 | -30, -46, 52 | 30, -43, 49 | -42, -4, 58 | 42, -4, 58 |
| Subject 9 | -51, -76, -2 | 51, -70, -2 | -33, -43, 55 | 30, -43, 49 | -42, -4, 58 | 42, -4, 58 |
| Subject 10 | -51, -76, -2 | 51, -70, -2 | -33, -43, 55 | 30, -43, 49 | -42, -4, 58 | 42, -4, 58 |
| Subject 11 | -51, -76, -2 | 51, -70, -2 | -30, -46, 52 | 30, -43, 49 | -42, -4, 58 | 42, -4, 58 |
| Subject 12 | -51, -76, -2 | 51, -70, -2 | -30, -43, 52 | 30, -43, 52 | -42, -4, 58 | 42, -4, 58 |
| Subject 13 | -51, -76, -2 | 51, -72, -2 | -30, -46, 52 | 30, -40, 49 | -42, -4, 58 | 42, -4, 58 |
| Subject 14 | -48, -67, 4 | 51, -70, -2 | -30, -43, 52 | 30, -43, 52 | -42, -4, 58 | 42, -4, 58 |
| Subject 15 | -51, -76, -2 | 51, -70, -2 | -30, -43, 52 | 30, -40, 46 | -42, -4, 58 | 42, -4, 58 |
| Subject 16 | -51, -76, -2 | 51, -70, -2 | -30, -46, 52 | 30, -43, 49 | -42, -4, 58 | 42, -4, 58 |
| Subject 17 | -51, -76, -2 | 51, -70, -2 | -30, -43, 52 | 30, -40, 46 | -42, -7, 58 | 42, -7, 58 |
| Subject 18 | -51, -76, -2 | 51, -70, -2 | -30, -43, 52 | 30, -43, 49 | -42, -4, 58 | 42, -4, 58 |
| Subject 20 | -51, -73, -2 | 51, -70, -2 | -30, -43, 52 | 30, -40, 49 | -42, -4, 58 | 42, -4, 58 |
| Subject 21 | -51, -76, -2 | 51, -70, -2 | -30, -43, 52 | 30, -43, 49 | -42, -4, 58 | 42, -4, 58 |
| Subject 22 | -51, -76, -2 | 51, -70, -2 | -30, -43, 52 | 30, -43, 52 | -42, -4, 58 | 42, -4, 58 |
| Subject 23 | -51, -76, -2 | 51, -70, -2 | -30, -43, 52 | 30, -43, 49 | -42, -4, 58 | 42, -4, 58 |
| Subject 24 | -48, -73, -2 | 51, -70, -2 | -30, -43, 52 | 30, -43, 52 | -42, -4, 58 | 42, -4, 58 |
| Subject 25 | -51, -76, -2 | 51, -70, -2 | -30, -43, 52 | 30, -43, 49 | -42, -4, 58 | 42, -4, 58 |
| Subject 26 | -51, -76, -2 | 51, -70, -2 | -30, -43, 52 | 30, -40, 46 | -42, -4, 58 | 42, -4, 58 |
| Subject 27 | -51, -76, -2 | 51, -70, -2 | -30, -46, 52 | 30, -43, 52 | -42, -4, 58 | 42, -4, 58 |
| Subject 28 | -51, -76, -2 | 51, -70, -2 | -30, -43, 52 | 30, -43, 52 | -42, -4, 58 | 42, -4, 58 |
| Subject 29 | -51, -76, -2 | 51, -70, -2 | -30, -46, 52 | 30, -43, 49 | -42, -4, 58 | 42, -4, 58 |
| Subject 30 | -51, -76, -2 | 51, -70, -2 | -30, -43, 52 | 30, -43, 52 | -42, -4, 58 | 42, -4, 58 |

**Figure S1.**
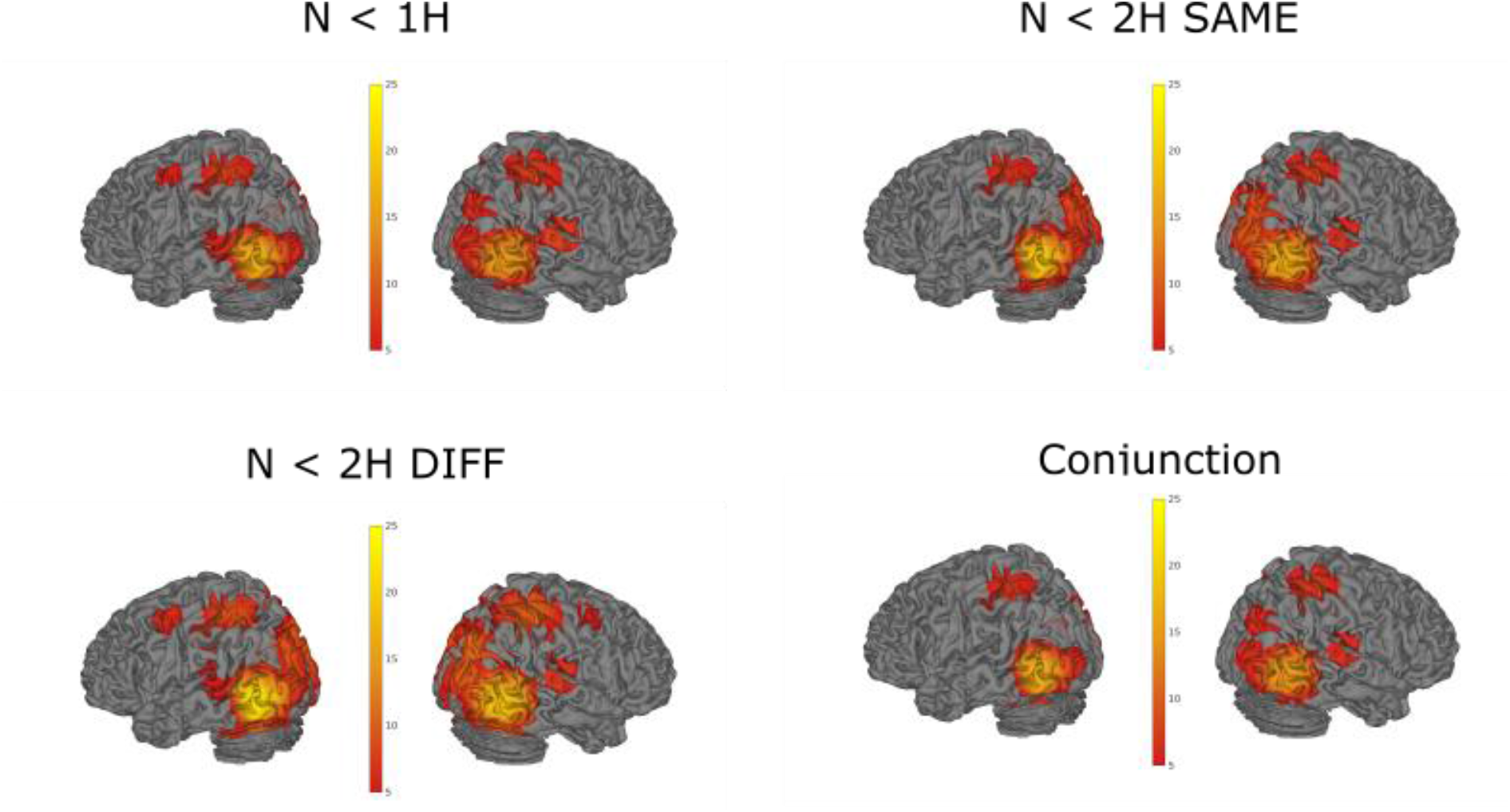
Brain activation in each of the action observation conditions compared with the neutral condition. The bottom right figure shows the conjunction of the other three contrasts (Nichols et al. 2005). Images were thresholded using a *p* < .05 FWE-corrected threshold. Left premotor activation was also present in the conjunction analysis at an uncorrected threshold of *p* < .001.

**Figure S2.**
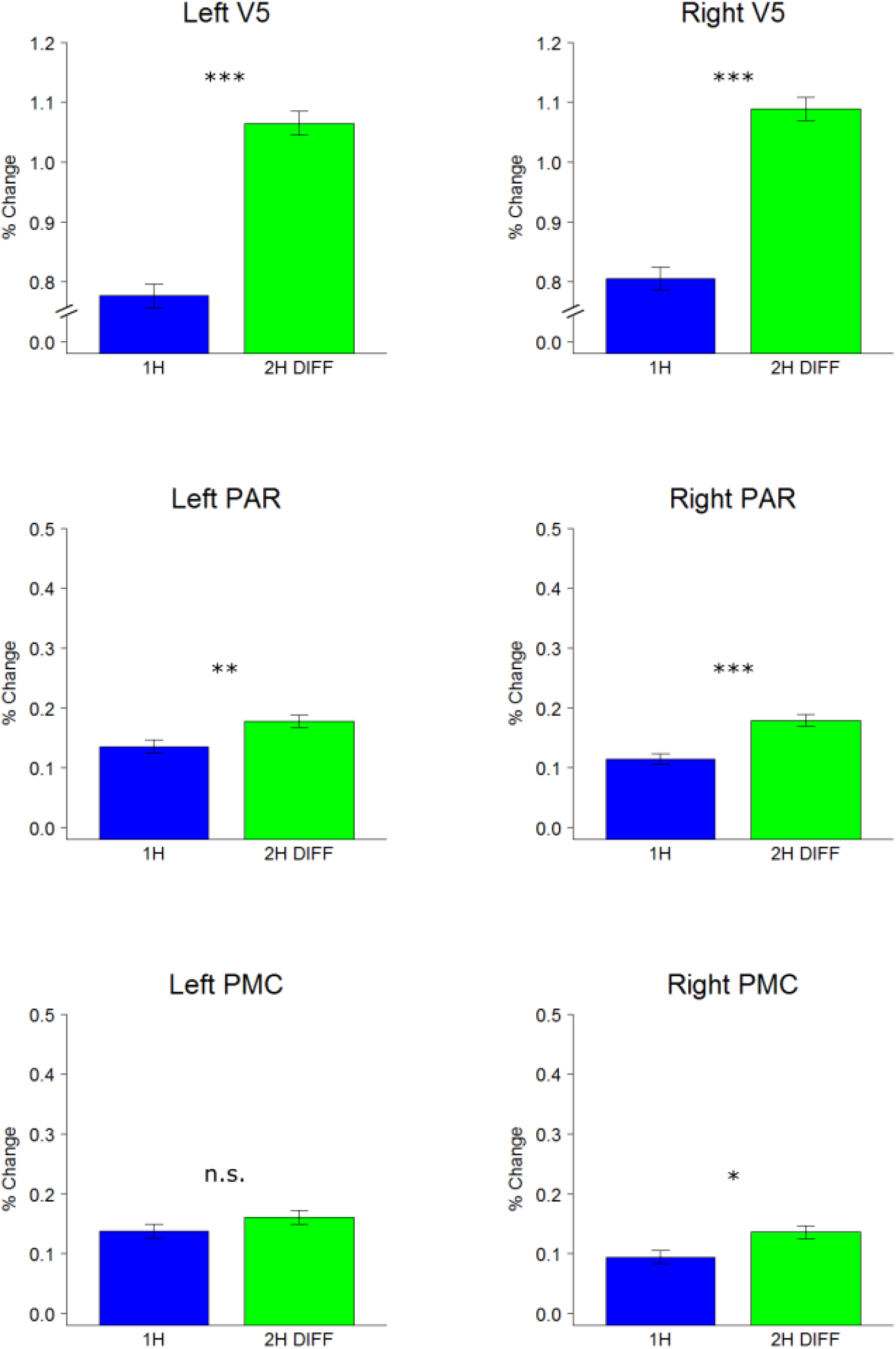
Hemisphere specific results of the ROI analyses testing whether activation was stronger in the 2H DIFF condition than in the 1H condition. The y-axis shows the % signal change with respect to the N condition. Details on the % signal change calculation are provided in the methods. Post-hoc two-tailed *t* tests comparing 1H with 2H DIFF are displayed. Error bars are SEMs corrected for within-subject designs (Morey 2008). * p < .05; ** p < .01; *** p < .001.

**Figure S3.**
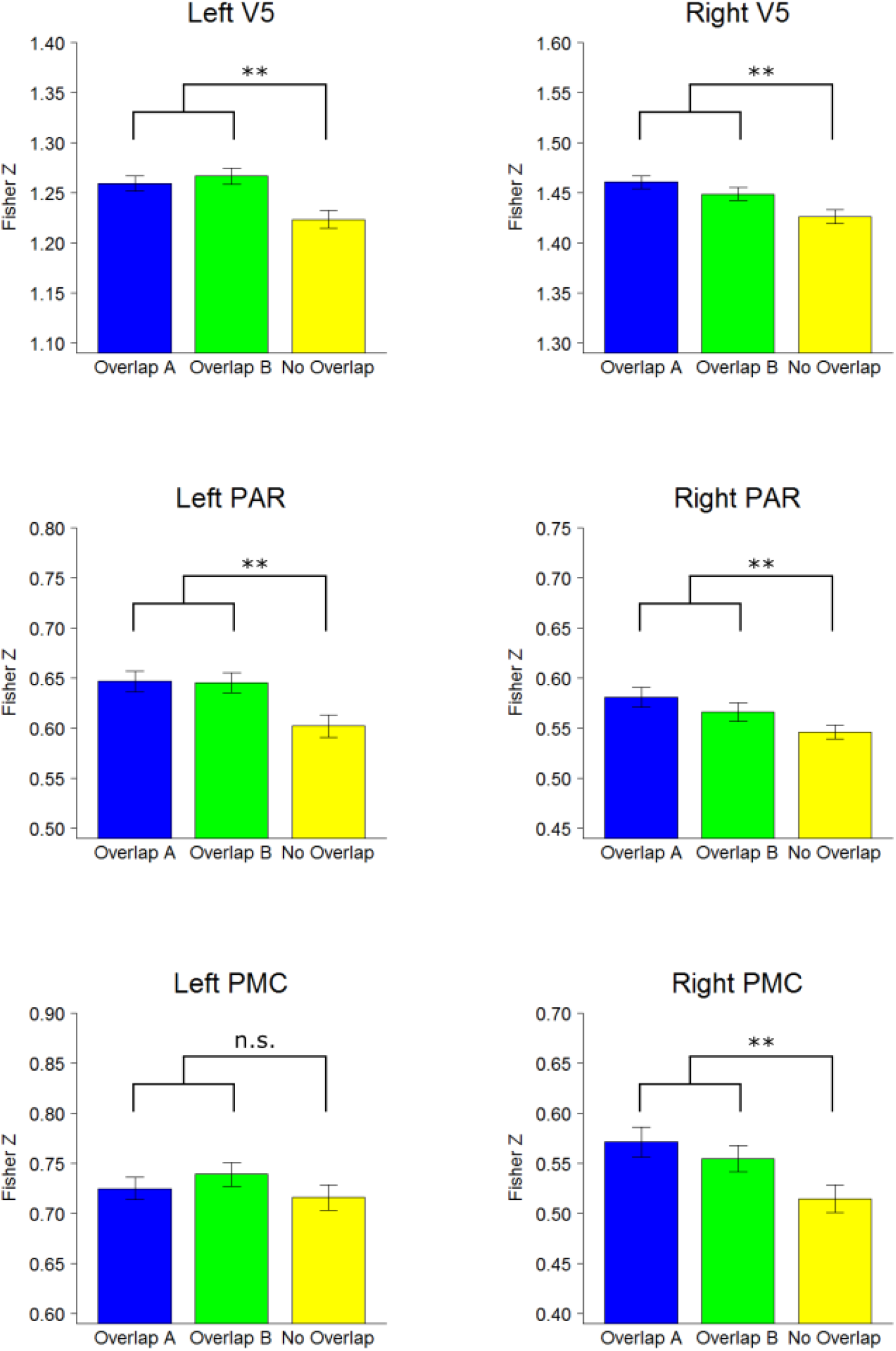
Hemisphere specific results of the representational similarity analyses testing whether the two observed gestures in the 2H DIFF condition could be decoded from brain activation in the three ROIs. Post-hoc two-tailed *t* tests comparing the average of Overlap A and B with No Overlap are displayed. Error bars are SEMs corrected for within-subject designs (Morey 2008). The difference between Overlap A and Overlap B was never significant. * p < .05; ** p < .01; *** p < .001.

Video *S1*. Video showing the procedure of the action execution task.

Video *S2*. Video showing the 1H condition of the action observation task.

Video *S3*. Video showing the 2H SAME condition of the action observation task.

Video *S4*. Video showing the 2H DIFF condition of the action observation task. This video also illustrates the glitches participants had to detect in the videos.

## References

Aglioti SM, Cesari P, Romani M, Urgesi C. 2008. Action anticipation and motor resonance in elite basketball players. Nat Neurosci. 11:1109–1116.

Aihara T, Yamamoto S, Mori H, Kushiro K, Uehara S. 2015. Observation of interactive behavior increases corticospinal excitability in humans: A transcranial magnetic stimulation study. Brain Cogn. 100:1–6.

Arnstein D, Cui F, Keysers C, Maurits NM, Gazzola V. 2011. Suppression during Action Observation and Execution Correlates with BOLD in Dorsal Premotor, Inferior Parietal, and SI Cortices. J Neurosci. 31:14243–14249.

Avenanti A, Candidi M, Urgesi C. 2013. Vicarious motor activation during action perception: beyond correlational evidence. Front Hum Neurosci. 7:1–8.

Babiloni C, Del Percio C, Vecchio F, Sebastiano F, Di Gennaro G, Quarato PP, Morace R, Pavone L, Soricelli A, Noce G, Esposito V, Rossini PM, Gallese V, Mirabella G. 2016. Alpha, beta and gamma electrocorticographic rhythms in somatosensory, motor, premotor and prefrontal cortical areas differ in movement execution and observation in humans. Clin Neurophysiol. 127:641–654.

Botvinick MM, Braver TS, Barch DM, Carter CS, Cohen JD. 2001. Conflict Monitoring and Cognitive Control. Psychol Rev. 108:624–652.

Botvinick MM, Cohen JD, Carter CS. 2004. Conflict monitoring and anterior cingulate cortex: An update. Trends Cogn Sci. 8:539–546.

Brass M, Derrfuss J, von Cramon DY. 2005. The inhibition of imitative and overlearned responses: a functional double dissociation. Neuropsychologia. 43:89–98.

Brass M, Ruby P, Spengler S. 2009. Inhibition of imitative behaviour and social cognition. Philos Trans R Soc B-biological Sci. 364:2359–2367.

Braver TS, Barch DM, Gray JR, Molfese DL, Snyder A. 2001. Anterior cingulate cortex and response conflict: Effects of frequency, inhibition and errors. Cereb Cortex. 11:825–836.

Brett M, Anton JL, Valabregue R, Poline JB. 2002. Region of interest analysis using an SPM toolbox. Abstr Present 8th Int Conf Funct Mapp Hum Brain.

Bucchioni G, Cavallo A, Ippolito D, Marton G, Castiello U. 2013. Corticospinal excitability during the observation of social behavior. Brain Cogn. 81:176–182.

Caspers S, Zilles K, Laird AR, Eickhoff SB. 2010. ALE meta-analysis of action observation and imitation in the human brain. Neuroimage. 50:1148–1167.

Centelles L, Assaiante C, Nazarian B, Anton JL, Schmitz C. 2011. Recruitment of both the mirror and the mentalizing networks when observing social interactions depicted by point-lights: A neuroimaging study. PLoS One. 6.

Colling LJ, Knoblich GG, Sebanz N. 2013. How does “mirroring” support joint action? Cortex. 49:2964–2965.

Cracco E, Bardi L, Desmet C, Genschow O, Rigoni D, De Coster L, Radkova I, Deschrijver E, Brass M. 2018. Automatic Imitation: A meta-analysis. Psychol Bull. 144:453–500.

Cracco E, Brass M. 2018a. Automatic imitation of multiple agents: Simultaneous or random representation? J Exp Psychol Hum Percept Perform. 44:729–740.

Cracco E, Brass M. 2018b. The role of sensorimotor processes in social group contagion. Cogn Psychol. 103:23–41.

Cracco E, De Coster L, Andres M, Brass M. 2015. Motor simulation beyond the dyad: Automatic imitation of multiple actors. J Exp Psychol Hum Percept Perform. 41:1488–1501.

Cracco E, De Coster L, Andres M, Brass M. 2016. Mirroring multiple agents: Motor resonance during action observation is modulated by the number of agents. Soc Cogn Affect Neurosci. 11:1422–1427.

Desmet C, Deschrijver E, Brass M. 2014. How social is error observation? The neural mechanisms underlying the observation of human and machine errors. Soc Cogn Affect Neurosci. 9:427–435.

Esterman M, Tamber-Rosenau BJ, Chiu YC, Yantis S. 2010. Avoiding non-independence in fMRI data analysis: Leave one subject out. Neuroimage. 50:572–576.

Fadiga L, Fogassi L, Pavesi G, Rizzolatti G. 1995. Motor facilitation during action observation: A magnetic stimulation study. J Neurophysiol. 73:2608–2611.

Fouragnan E, Retzler C, Philiastides MG. 2018. Separate neural representations of prediction error valence and surprise: Evidence from an fMRI meta-analysis. Hum Brain Mapp. 2887–2906.

Fox NA, Bakermans-Kranenburg MJ, Yoo KH, Bowman LC, Cannon EN, Vanderwert RE, Ferrari PF, van IJzendoorn MH. 2016. Assessing human mirror activity with EEG mu rhythm: A meta-analysis. Psychol Bull. 142:291–313.

Gallese V, Fadiga L, Fogassi L, Rizzolatti G. 1996. Action recognition in the premotor cortex. Brain. 119:593–609.

Gazzola V, Keysers C. 2009. The observation and execution of actions share motor and somatosensory voxels in all tested subjects: Single-subject analyses of unsmoothed fMRI data. Cereb Cortex. 19:1239–1255.

Georgescu AL, Kuzmanovic B, Santos NS, Tepest R, Bente G, Tittgemeyer M, Vogeley K. 2014. Perceiving Nonverbal Behavior: Neural Correlates of Processing Movement Fluency and Contingency in Dyadic Interactions. Hum Brain Mapp. 35:1362–1378.

Hadley L V., Novembre G, Keller PE, Pickering MJ. 2015. Causal Role of Motor Simulation in Turn-Taking Behavior. J Neurosci. 35:16516–16520.

Hamilton AFDC, Grafton ST. 2008. Action outcomes are represented in human inferior frontoparietal cortex. Cereb Cortex. 18:1160–1168.

Hutzler F. 2014. Reverse inference is not a fallacy per se: Cognitive processes can be inferred from functional imaging data. Neuroimage. 84:1061–1069.

Iacoboni M, Lieberman MD, Knowlton BJ, Molnar-Szakacs I, Moritz M, Throop CJ, Fiske AP. 2004. Watching social interactions produces dorsomedial prefrontal and medial parietal BOLD fMRI signal increases compared to a resting baseline. Neuroimage. 21:1167–1173.

Keysers C, Gazzola V. 2006. Towards a unifying neural theory of social cognition. In:Anders,, Ende,, Junghöfer,, Kissler,, Wildgruber, editors. Progress in Brain Research. Elsevier B.V. p. 379–401.

Kilner JM, Lemon RN. 2013. What we know currently about mirror neurons. Curr Biol. 23:R1057–R1062.

Kourtis D, Sebanz N, Knoblich G. 2013. Predictive representation of other people’s actions in joint action planning: An EEG study. Soc Neurosci. 8:31–42.

Kriegeskorte N, Mur M, Bandettini P. 2008. Representational similarity analysis – connecting the branches of systems neuroscience. Front Syst Neurosci. 2:1–28.

Liston C, Matalon S, Hare TA, Davidson MC, Casey BJ. 2006. Anterior Cingulate and Posterior Parietal Cortices Are Sensitive to Dissociable Forms of Conflict in a Task-Switching Paradigm. Neuron. 50:643–653.

Liu X, Banich MT, Jacobson BL, Tanabe JL. 2004. Common and distinct neural substrates of attentional control in an integrated Simon and spatial Stroop task as assessed by event-related fMRI. Neuroimage. 22:1097–1106.

Mayer AR, Teshiba TM, Franco AR, Ling J, Shane MS, Stephen JM, Jung RE. 2012. Modeling conflict and error in the medial frontal cortex. Hum Brain Mapp. 33:2843–2855.

McLaren DG, Ries ML, Xu G, Johnson SC. 2012. A generalized form of context-dependent psychophysiological interactions (gPPI): A comparison to standard approaches. Neuroimage. 61:1277–1286.

Milham MP, Banich MT, Webb A, Barad V, Cohen NJ, Wszalek T, Kramer AF. 2001. The relative involvement of anterior cingulate and prefrontal cortex in attentional control depends on nature of conflict. Cogn Brain Res. 12:467–473.

Molenberghs P, Cunnington R, Mattingley JB. 2012. Brain regions with mirror properties: A meta-analysis of 125 human fMRI studies. Neurosci Biobehav Rev. 36:341–349.

Morey RD. 2008. Confidence intervals from normalized data: A correction to Cousineau (2005). Tutor Quant Methods Psychol. 4:61–64.

Mukamel R, Ekstrom AD, Kaplan JT, Iacoboni M, Fried I. 2010. Single-Neuron Responses in Humans during Execution and Observation of Actions. Curr Biol. 20:750–756.

Naish KR, Houston-Price C, Bremner AJ, Holmes NP. 2014. Effects of action observation on corticospinal excitability: Muscle specificity, direction, and timing of the mirror response. Neuropsychologia. 64:331–348.

Nichols T, Brett M, Andersson J, Wager T, Poline JB. 2005. Valid conjunction inference with the minimum statistic. Neuroimage. 25:653–660.

Oldfield RC. 1971. Assessment and analysis of handedness – Edinburgh Inventory. Neuropsychologia. 9:97–113.

Poldrack RA. 2006. Can cognitive processes be inferred from neuroimaging data? Trends Cogn Sci. 10:59–63.

Quadflieg S, Koldewyn K. 2017. The neuroscience of people watching: How the human brain makes sense of other people’s encounters. Ann N Y Acad Sci. 1396:166–182.

Quadflieg S, Penton-Voak IS. 2017. The Emerging Science of People-Watching: Forming Impressions From Third-Party Encounters. Curr Dir Psychol Sci. 26:383–389.

Ridderinkhof KR, Ullsperger M, Crone E a, Nieuwenhuis S. 2004. The role of the medial frontal cortex in cognitive control. Science (80- ). 306:443–447.

Rizzolatti G, Sinigaglia C. 2010. The functional role of the parieto-frontal mirror circuit: Interpretations and misinterpretations. Nat Rev Neurosci. 11:264–274.

Rizzolatti G, Sinigaglia C. 2016. The mirror mechanism: a basic principle of brain function. Nat Rev Neurosci. 17:757–765.

Tsai JCC, Sebanz N, Knoblich GG. 2011. The GROOP effect: Groups mimic group actions. Cognition. 118:135–140.

Ullsperger M, Danielmeier C, Jocham G. 2014. Neurophysiology of performance monitoring and adaptive behavior. Physiol Rev. 94:35–79.

Urgesi C, Candidi M, Avenanti A. 2014. Neuroanatonnical substrates of action perception and understanding: an anatomic likelihood estimation meta-analysis of lesion-symptom mapping studies in brain injured patients. Front Hum Neurosci. 8:344.

van Veen V, Cohen JD, Botvinick MM, Stenger VA, Carter CS. 2001. Anterior Cingulate Cortex, Conflict Monitoring, and Levels of Processing. Neuroimage. 14:1302–1308.

Verbruggen F, Notebaert W, Liefooghe B, Vandierendonck A. 2006. Stimulus- and response-conflict-induced cognitive control in the flanker task. Psychon Bull Rev. 13:328–333.

Volpe G, D’Ausilio A, Badino L, Camurri A, Fadiga L. 2016. Measuring social interaction in music ensembles Measuring social interaction in music ensembles. Philos Trans R Soc B Biol Sci. 371:20150377.

Wendelken C, Ditterich J, Bunge S a, Carter CS. 2009. Stimulus and response conflict processing during perceptual decision making. Cogn Affect Behav Neurosci. 9:434–447.

Yarkoni T, Poldrack RA, Nichols TE, Van Essen DC, Wager TD. 2011. Large-scale automated synthesis of human functional neuroimaging data. Nat Methods. 8:665–670.

